# Discovery of a high-performance phage-derived promoter/repressor system for probiotic lactobacillus engineering

**DOI:** 10.1101/2023.12.05.570180

**Authors:** Marc Blanch-Asensio, Varun Sai Tadimarri, Alina Wilk, Shrikrishnan Sankaran

## Abstract

**Background:** The *Lactobacillus* family comprises many species of great importance for the food and healthcare industries, with numerous strains identified as beneficial for humans and used as probiotics. Hence, there is a growing interest in engineering these probiotic bacteria as live biotherapeutics for animals and humans. However, the genetic parts needed to regulate gene expression in these bacteria remain limited compared to model bacteria like *E. coli* or *B. subtilis*. To address this deficit, in this study, we selected and tested several bacteriophage-derived genetic parts with the potential to regulate transcription in lactobacilli.

**Results:** We screened genetic parts from 6 different lactobacilli-infecting phages and identified one promoter/repressor system with unprecedented functionality in *L. plantarum* WCFS1. The phage-derived promoter was found to achieve expression levels nearly 9-fold higher than the previously reported strongest promoter in this strain and the repressor was able to almost completely repress this expression by reducing it nearly 500-fold.

**Conclusions:** The new parts and insights gained from their engineering will enhance the genetic programmability of lactobacilli for healthcare and industrial applications.

## BACKGROUND

Lactobacilli are ubiquitous in and around humans, providing numerous health benefits. They are commensals in multiple organs (gut, skin, urinary tract, vagina, lungs, etc.), and are one of the largest families of probiotics [1]. Several probiotic strains are being clinically tested as Live Biotherapeutic Products to treat diseases like ulcerative colitis [2], mastitis [3], atopic dermatitis [4], bacterial vaginosis [5], and periodontitis [6]. Apart from this, lactobacilli are also vital in the food industry for producing fermented products like yogurt [7], cheese [8], beer [9], wine [10]. etc. Due to this close association with our lives, there is considerable interest to genetically enhance and expand their capabilities for both medical [11] and industrial applications [12]. In the medical sphere, lactobacilli are being engineered as drug delivery vehicles to treat diseases like ulcerative colitis [13], HIV infection [14], respiratory infections [15], etc. or as oral vaccine candidates that display antigens on their cell surface [16]. In the industrial sphere, lactobacilli are being considered as alternative recombinant expression hosts to *E. coli* since (i) they don’t produce endotoxins and many strains have Generally Recognized as Safe (GRAS) status, minimizing the toxicity risks for pharmaceutical protein production [17], (ii) infrastructure for their culture is well established in the food industry, and (iii) they are good at secreting proteins that can be purified from culture supernatants [18].

To realize such applications, considerable progress has been made in identifying a handful of genetic parts including constitutive promoters [19], peptide-inducible promoters [20], ribosome binding sites (RBS) [21], signal peptides for protein secretion [18], origins of replication [22], and food-grade plasmid retention systems [23, 24]. However, the programmability of lactobacilli is severely limited by the scarcity of genetic parts that allow control over gene expression, especially strong promoters and effective repressors that are needed to realize important functions like inducible expression.

Synthetic biology tools enable us to genetically program microorganisms with functions analogous to electronic circuits [25, 26]. The most basic element for such programming is a switch that can regulate the expression of genes between ON and OFF states depending on the presence and absence of a trigger. These switches are typically inducible gene expression systems, with which the production of recombinant proteins can be activated by an inducer molecule (e.g., IPTG) [27] or stimulus (e.g., light) [28]. These switches most often involve repressors, whose function is to block gene expression driven by a promoter [29]. While such screening and identification of repressors has been extensively done in model organisms like *Escherichia coli* [30, 31], lactobacilli severely lack reliable repressors.

Heiss and colleagues performed a study in *L. plantarum* in which they tested several promoter/repressor systems either derived from *Bacillus megaterium*, *E. coli* or other lactic acid bacteria (LAB) but were found to be considerably leaky and the promoters associated with them were weak [32]. For example, an endogenous promoter/repressor (*P_lacA_*/lacR) system showed a ∼8-fold induction capability in response to 2% (w/v) lactose in *L. plantarum* 3NSH. The system was completely repressed in the presence of monomeric sugars (glucose and galactose). In the same study, the orthogonal xylose inducible promoter/repressor system (*P_xylA_*_/_xylR), derived from *B. megaterium*, was responsive to xylose supplementation but showed significant leaky expression in the absence of the inducer. This study also tested the promoter/repressor system *P_lacA_*/lacI from *E. coli* in combination with T7 RNA Polymerase system. Post induction, the dynamic range of the system accounted for a ∼6-fold higher reporter expression. However, the reporter expression levels were low compared to a moderately strong constitute promoter. Overall, these results highlight the need for identifying promoter/repression systems that work reliably in *L. plantarum* (e.g., a strong repressor able to fully repress a strong promoter). In this study, we looked for repressors in another promising source for genetic parts – bacteriophages that infect LABs. By screening through multiple phage-derived repressors that control their lytic and lysogenic cycles in LABs, for the first time, we identified one candidate that efficiently and reliably represses gene expression in *L. plantarum* WCFS1. Interestingly, the native promoter associated with the repressor was found to drive the highest reported levels of gene expression in *L. plantarum* WCFS1. The discovery of this promoter and repressor combination lays the foundation for creating inducible gene expression systems and achieving advanced programming capabilities in lactobacilli.

## MATERIALS AND METHODS

### Strain, media and plasmids

*L*. *plantarum* WCFS1 was used as the parent strain in this study. The strain was maintained in the De Man, Rogosa and Sharpe (MRS) media (Carl Roth GmbH, Germany, Art. No. X924.1). Genetically engineered *L*. *plantarum* WCFS1 strains were grown in MRS media supplemented with 10 μg/mL of erythromycin (Carl Roth GmbH, Art. No. 4166.2) at 37°C and 250 revolutions per minute (rpm) for approximately 16 h. NEB 5-alpha Competent *E*. *coli* cells were used (New England Biolabs GmbH, Germany, Art. No. C2987) for the cloning of certain plasmids. This strain was maintained in Luria-Bertani (LB) media (Carl Roth GmbH, Art. No. X968.1). Genetically engineered *E*. *coli* DH5α strains were grown in LB media supplemented with 200 μg/mL of erythromycin at 37°C, 250 rpm shaking conditions for approximately 16 h. The pLp_3050sNuc plasmid, which was used as the backbone vector in this study, was a kind gift from Prof. Geir Mathiesen (Addgene plasmid # 122030). *E. coli* Nissle was a kind gift from Prof. Rolf Müller. *L. plantarum* Lg1e was a kind gift from Dr. Makiko Kakikawa.

### Molecular biology

Polymerase Chain Reaction (PCR) was performed using Q5 High Fidelity 2X Master Mix (NEB) with primers synthesized by Integrated DNA Technologies (IDT) (Leuven, Belgium) or Eurofins Genomics GmbH (Köln, Germany). Primers are listed in **Table S2**. Synthetic genes were purchased as eBlocks from IDT (Coralville, USA). The eBlocks were codon optimized using the IDT Codon Optimization Tool (Coralville, USA). NEBuilder® HiFi DNA Assembly Cloning Kit, Quick Blunting Kit and the T4 DNA Ligase enzyme were purchased from New England BioLabs (NEB, Germany). The plasmid extraction kit was purchased from Qiagen GmbH (Hilden, Germany). The DNA purification kit was purchased from Promega GmbH (Walldorf, Germany). Generuler 1 Kb DNA Ladder (Thermo Fisher Scientific) was used as a reference for the agarose gels.

### *L. plantarum* WCFS1 competent cell preparation and DNA transformation

*L. plantarum* WCFS1 was inoculated in 5 mL of MRS media without any antibiotic and grown overnight at 37 °C with shaking (250 rpm). The next day, 1 mL of the bacterial culture was transferred into a secondary culture based on 20 mL of MRS and 5 mL of 1% (w/v) glycine. The secondary culture was incubated at 37 °C, 250 rpm until the optical density of the sample measured at a wavelength of 600 nm (OD_600_) reached approximately 1. The cells were pelleted down by centrifuging at 4000 rpm for 12 min at 4°C. Next, the bacterial pellet was washed several times, in each, bacteria were centrifugated for 8 minutes at 4000 rpm. The first two washes were done with 5 mL of ice-cold 10 mM MgCl_2._ The next two washes were performed with 5 mL of ice-cold Sac/Gly solution [10% (v/v) glycerol and 1 M sucrose mixed in a 1:1 (v/v) ratio]. Lastly, after discarding the supernatant, the pellet was resuspended in 500 μL of Sac/Gly solution, and the competent cells were distributed in 60 μL aliquots for DNA transformation. For transformation, 1 μg of dsDNA was added to the competent cells and incubated on ice for 10 minutes. The mixture was transferred to an ice-cold 2 mm gap electroporation cuvette (Bio-Rad Laboratories GmbH, Germany), and cells were electroporated with a single pulse at 1.8 kV, after which 1 mL of MRS medium was immediately added. The mixture was then incubated at 37 °C, 250 rpm for a recovery period of 3 h. After the recovery, the cells were centrifuged at 4000 rpm for 5 min, and 800 μL of the supernatant was discarded. The remaining 200 μL were used to resuspend the pellet, and the entire 200 μL were plated on MRS Agar supplemented with 10 μg/mL of erythromycin. The plates were incubated at 37 °C for 1-3 days to allow the growth of bacterial colonies.

### Direct cloning in *L. plantarum* WCFS1

Plasmid engineering of *L. plantarum* WCFS1 was done using the direct cloning method previously developed by us [33], which involved PCR-based amplification and circularization of recombinant plasmids, which were then transformed in the bacteria by electroporation. In brief, complementary overhangs for HiFi Assembly were either synthesized as custom-designed eBlocks or generated by PCR. The HiFi DNA Assembly reaction was performed following the manufacturer’s protocol. Then, 5 μL of the assembled HiFi product was used as a DNA template in the PCR reaction (100 μL final volume). After purifying the PCR product, 1000 to 2000 ng of linear DNA was phosphorylated using the Quick Blunting Kit and following the manufacturer’s protocol. Next, phosphorylated products were ligated using the T4 ligase enzyme. Two ligation reactions were set per cloning, each based on 500 ng of phosphorylated DNA, 2.5 µl of 10X T4 Ligase Buffer and 1.5 µl of T4 Ligase enzyme (autoclaved Milli-Q water was added to make the reaction volume to 25 µl). The ligations were incubated at 25 °C for 3 to 5 hours and then at 70 °C for 30 min for enzyme inactivation. After the incubation, the ligations were mixed and purified, performing three rounds of elution to concentrate the DNA (each with 10 µl of autoclaved Milli-Q water). The entire eluted mix (approximately 1000 ng) was transformed into *L. plantarum* WCFS1 electrocompetent cells.

The same strategy was used to achieve site-directed mutagenesis, where specific DNA sequences were removed from the plasmid by PCR using the bacterial pellet as a template for the PCR and primers covering the whole region but the targeted sequence. The linear PCR product was circularized as described before using the Quick Blunting Kit and the T4 Ligase. For sequence verification, DNA sequences of interest were amplified (100 μL final volume) using a bacterial pellet as a template. The PCR product was purified and sent for Sanger sequencing to Eurofins Genomics GmbH (Köln, Germany) by selecting the additional DNA purification step prior to sequencing.

### *E. coli* Nissle 1917 competent cell preparation

Wild-type *E. coli* Nissle 1917 bacteria was grown overnight in in LB media at 37 °C, 250 rpm. The next day, bacteria were subcultured in 100 mL of fresh LB media and incubated at 37 °C and 250 rpm until the OD_600_ reached 0.4. Bacteria were pelleted down by centrifugation at 4000 rpm for 5 minutes. After discarding the supernatant, the pellet was washed twice with 10 mL of ice-cold CaCl2 (200 mM) and once with a 10 mL of 1:1 combination of CaCl2 (200 mM) and glycerol (10% w/v). Following the final wash, the pellet was resuspended in 1 mL of CaCl2 + glycerol mixture and 100 μL aliquots were prepared and stored at -80°C unless used immediately.

### *E. coli DH*5α and *E. coli* Nissle 1917 DNA transformation

*E. coli* DH5α DNA transformation was performed following the manufacturer’s protocol for the NEBuilder® HiFi DNA Assembly Cloning Kit. For *E. coli* Nissle 1917 DNA transformation, 200 ng of plasmid DNA were mixed well with the competent cells by pipetting gently and incubated on ice for 30 minutes. Following the incubation, a 45-second heat shock was performed by placing the cells at a 42°C water bath. Next, cells were again incubated on ice for 5 minutes. After that, 900 μl of SOC media was added to the cell mixture and kept for incubation for 1 hour at 37°C. Next, the mixture was pelleted down by centrifugation at 4000 rpm for 5 minutes. 600 μl were immediately discarded, and the remaining 300 μl were used to resuspend the mixture. Finally, 150 μl were plated on an LB agar plate supplemented with 200 μg/mL of erythromycin and incubated at 37°C overnight.

### Flow cytometry analysis

Engineered strains were grown in 5 mL of MRS media supplemented with 10 µg/mL erythromycin at 37 °C with shaking, 250 rpm. The next day, bacteria were subcultured to an OD_600_ of 0.01 in 5 mL of MRS media (supplemented with 10 µg/mL erythromycin) and grown at 37 °C with shaking (250 rpm) for 16h. The following day, 1 mL of the bacterial suspensions were harvested by centrifugation at 10000 rpm. After discarding the supernatant carefully, the pellet was resuspended in 1 mL of sterile Dulbecco’s 1X PBS. The mixtures were then serially diluted by a 10^4^ Dilution Factor, and 5,000 bacteria events were recorded for analysis using Guava easyCyte BG flow-cytometer (Luminex, USA). A predesigned gate based on forward side scatter and side scatter thresholding was employed to get rid of debris and doublets during the collection of events. The fluorescence intensity of mCherry was measured using excitation by a green laser at 532 nm (100 mW) and the Orange-G detection channel 620/52 nm filter was used for signal analysis. The gain settings used for the data recording were as follows: Forward Scatter (FSC) – 11.8, Side Scatter (SSC) - 4, and Orange-G Fluorescence – 1.68. The compensation control for fluorescence recording was set at 0.01, with an acquisition rate of 5 decades. The Luminex GuavaSoft 4.0 software for EasyCyte was used for the analysis and representation of data.

### Microplate reader setup for reporter gene expression quantification

*L. plantarum* WCSF1 engineered strains were grown in the same manner as described for the Flow Cytometry Analysis. 200 µL of the 1000-µL resuspended mixture (PBS containing engineered bacteria) was added to a UV STAR Flat Bottom 96 well microtiter plate (Greiner BioOne GmbH, Germany). Next, the samples were analyzed in the Microplate Reader Infinite 200 Pro (Tecan Deutschland GmbH, Germany) and both the absorbance (600 nm wavelength), and mCherry fluorescence intensity (Ex_λ_ / Em_λ_ = 587 nm/625 nm) were measured. The Z-position and gain settings were set to 19000 µm and 100, respectively. The readings were taken using the top read setting. The fluorescence values were normalized with the optical density of the bacterial cells to calculate the Relative Fluorescence Units (RFU) (formula RFU = Fluorescence/OD_600_). The same procedure was followed for *E. coli* Nissle engineered strains, but those were grown in LB media supplemented with 200 μg/mL of erythromycin at 37°C with shaking (250 rpm). Experiments were performed in triplicates on three different days.

### Growth rate measurements and biomass calculation

To measure the growth curves of the engineered strains, they were cultivated overnight in antibiotic supplemented MRS media at 37°C with shaking (250 rpm). The following day, the bacterial cultures were subcultured in 3-mL secondary cultures at an initial OD_600_ = 0.01 and incubated at 30°C with shaking (250 rpm) until reaching an OD_600_ of 0.4-0.5. Then, 200 µL of the cultures were distributed in a UV STAR Flat Bottom 96 well microtiter plate. The plate was placed in the Microplate Reader with constant shaking conditions at an incubation temperature of 37°C. The kinetic assay was set to record the absorbance (600 nm) of the bacterial cultures with an interval of 15 min for 16 hours. The experiment was conducted twice on two independent days, keeping two technical duplicates per experiment.

To estimate the bacterial biomass of the engineered strains, they were cultivated overnight in antibiotic supplemented MRS media at 37°C with shaking (250 rpm). The following day, the bacterial cultures were subcultured in 5-mL secondary cultures at an initial OD_600_ = 0.01 and incubated at 37°C with shaking (250 rpm) for 16h. The following day, all 5-mL cultures were pelleted down in several rounds of centrifugation (10000 rpm), and the biomass of the bacterial pellets was measured using an analytical balance (Denver Instrument). Experiments were performed in triplicates on three different days.

### Fluorescence microscopy analysis

Engineered strains were grown in the same manner as described for the Flow Cytometry Analysis. 10 µL of the 1000-µL resuspended mixture (PBS containing engineered bacteria) was placed on glass slides of 1.5Lmm thickness (Paul Marienfeld GmbH, Germany) and 1.5H glass coverslips (Carl Roth GmbH, Germany) were placed on top of it. The samples were then observed under the Plan Apochromat 100× oil immersion lens (BZ-PA100, NA 1.45, WD 0.13Lmm) of the Fluorescence Microscope BZ-X800 (Keyence Corporation, Illinois, USA). The mCherry signal was captured in the BZ-X TRITC filter (model OP-87764) at an excitation wavelength of 545/25Lnm and an emission wavelength of 605/70Lnm with a dichroic mirror wavelength of 565Lnm. The images were adjusted for identical brightness and contrast settings. ImageJ2 software was used to process the images.

### Statistical and bioinformatics analysis

Statistical analysis was performed using GraphPad Prism 7.0 software. Student’s T-tests were used to determine if there were significant differences between the means of the groups. InterPro was used to identify the DBD of rep. AlphaFold was used to predict the 3D structures of the rep repressor.

## RESULTS

### Strategy to identify reliable promoter/repressor systems

A strategy to find and optimize reliable promoter/repressor systems in *L. plantarum* WCFS1 was developed as shown in **Figure 1**. The search for potential transcriptional repressors was limited to repressors encoded in the genetic switches that regulate lytic and lysogenic cycles in bacteriophages. The strategy involved 1) identifying bacteriophages that infect lactobacilli with characterized genetic switches, 2) selecting 6 repressors with known operator sequences, 3) designing all the genetic parts required for 4) building a genetic platform, 5) testing the repression mediated by each repressor, and finally 6) improving such repression by introducing certain modifications to the operator/promoter regions.

**Figure 1:**
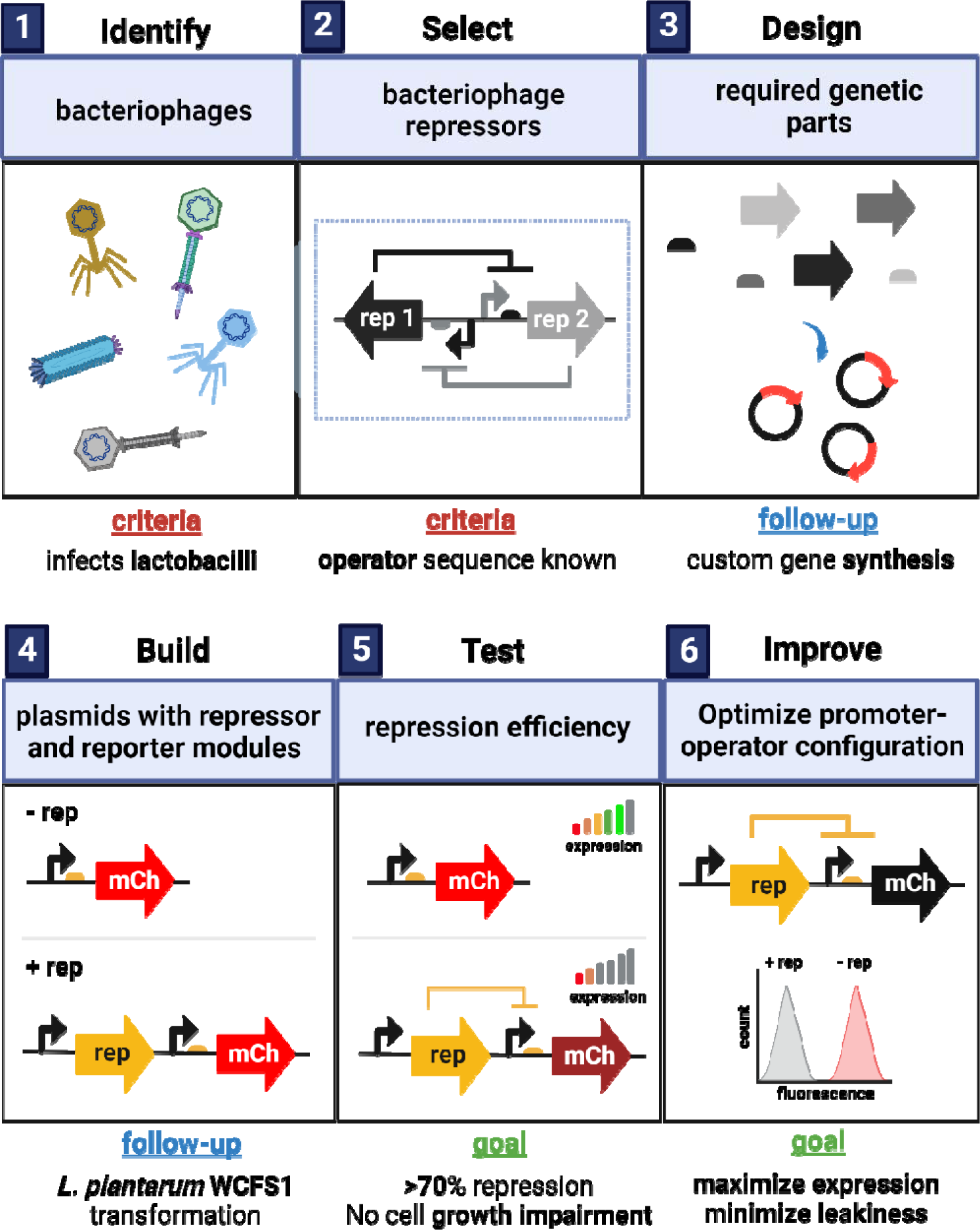
Panels showing the strategy and all the steps that were defined and followed to find novel bacteriophage-derived promoter/repressor systems in *L. plantarum* WCFS1.

### Screening of six bacteriophage repressors

The six different phage-derived repressors that we tested in this study are: <u>cng</u> and <u>cpg</u> repressors from the Lg1e phage infecting *L. plantarum* [34, 35], tec and rep repressors from the mv4 phage infecting *Lactobacillus delbrueckii* [36], and <u>cI</u> and <u>cro</u> repressors from the A2 phage infecting *Lactobacillus casei* [37, 38] (**Table S2).** A standard, simple genetic module was designed and built to test the repression mediated by each repressor. The module was based on our previously reported strongest constitutive promoter in *L. plantarum* WCFS1 (*P_tlpA_*) [39], which enabled reliable characterization of repressor activity. Notably, previous studies attempting to identify efficient repressors in lactobacilli were limited by promoters driving moderate levels of expression [32]. Our repressor-testing module included (i) *P_tlpA_*driving the expression of the reporter gene mCherry, (ii) repressor-specific operators inserted between the -10 box and the RBS (**Figure S1**) and (iii) repressors constitutively expressed by a moderately strong promoter (*P_48_*). The module was constructed through two rounds of cloning to encode both the operator and repressor in the plasmid. In the first round, the operator was inserted within the promoter, and in the second round, each repressor was cloned in the plasmid containing the corresponding operator (**Figure 2A**). Repression of mCherry production was first assessed by quantifying the drop in fluorescence intensity using flow cytometry (**Figure 2C**). This analysis first revealed that insertion of the operator sequences weakened the strength of *P_tlpA_* in all cases between ∼1.3 and 4.3-fold (**Figure S2**), although fluorescence intensities remained high enough to assess repressor activity. When the repressors were encoded in the plasmids containing the operator sequences, drop in fluorescence was observed only with the repressors cng, rep and cI (**Figure 2C**). Notably, rep was found to be the strongest repressor among the three with fluorescence intensity values comparable to the wild-type strain that was not modified to produce mCherry (**Figure S3**).

**Figure 2:**
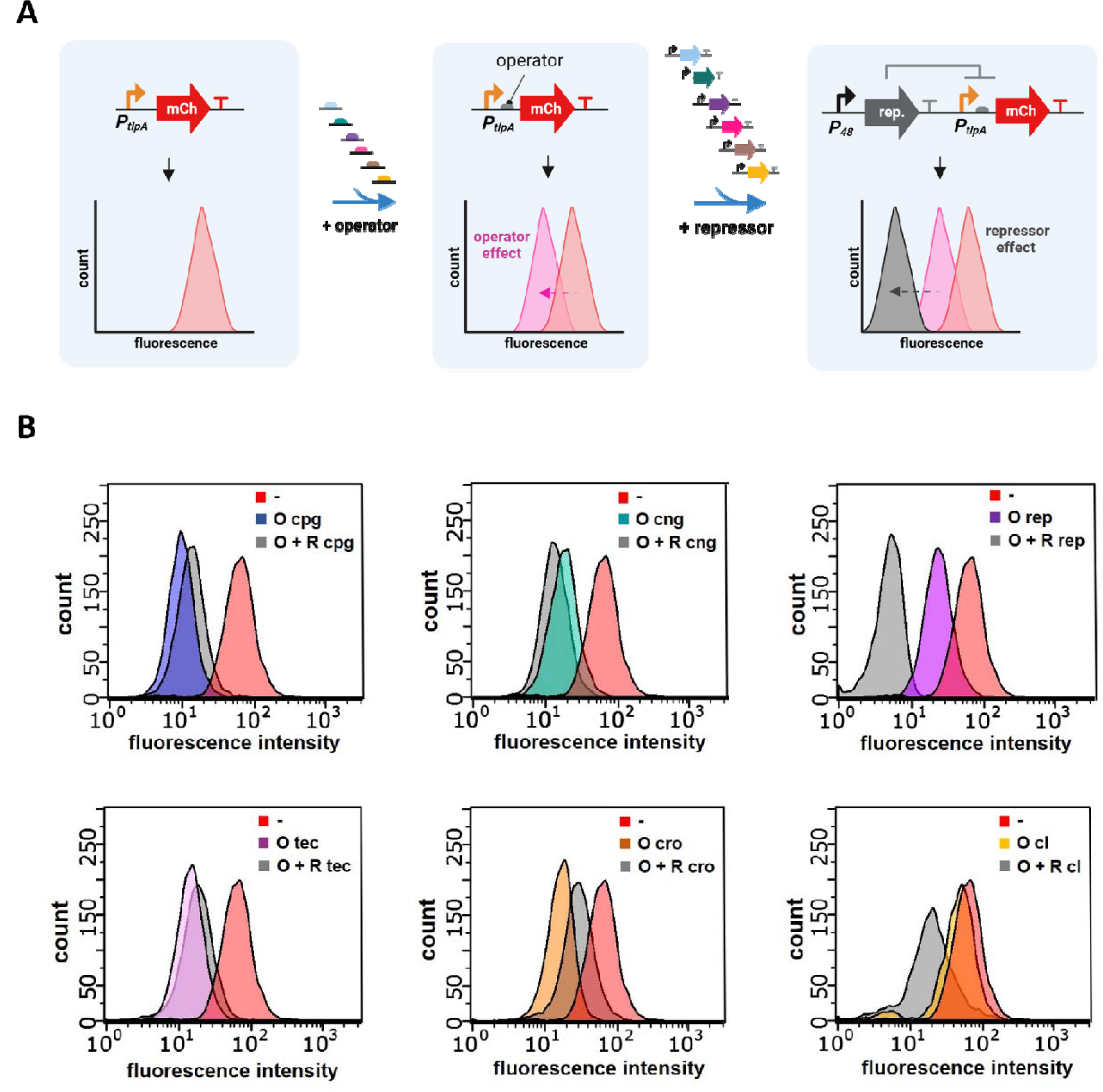
**A)** Cloning workflow design. In the first round of cloning, the operators were inserted within the promoter *P_tlpA_*. In the second round, the repressors were cloned in the corresponding plasmid. All genetic fragments were based on IDT synthetic eBlocks. **B)** Flow cytometry data showing the effect on mCherry expression of cloning each operator (O) and each operator plus repressor (O + R) in the plasmid.

### Quantification of repression levels

Next, we proceeded to quantify the repression levels and the effect on cell growth imposed by cng, rep and cI. We set a couple of requirements that potential repressors should meet in order to proceed with the next step of optimizing repression. Repressors i) should be able to repress at least up to 70% mCherry expression and ii) do not considerably impair bacterial growth. Concerning repression, only rep showed levels of repression above 70%, precisely, rep-driven repression was found to be up to 92% (**Figure 3A, 3B**). As for the effect on cell growth, we measured the growth curves of bacteria encoded with all three repressors in a microplate reader and compared them to wild-type bacteria (**Figure 3C**). Only the growth curve of the cI repressor was noticeably lower than the others. Nevertheless, differences in the OD_600_ after 16 hours of growth in the micro plate reader were found to be non-significant (**Figure S4**). In addition, we calculated the bacterial biomass after overnight growth in an incubator at 37°C with continuous shaking. In this scenario, bacteria constitutively expressing the cI repressor grew very poorly (∼85% lower biomass than WT), while rep and cng clones grew similar to bacteria carrying the *PtlpA*_mCherry plasmid (∼50% drop in biomass compared to WT) (**Figure 3D**). Overall, only the rep repressor met both requirements.

**Figure 3:**
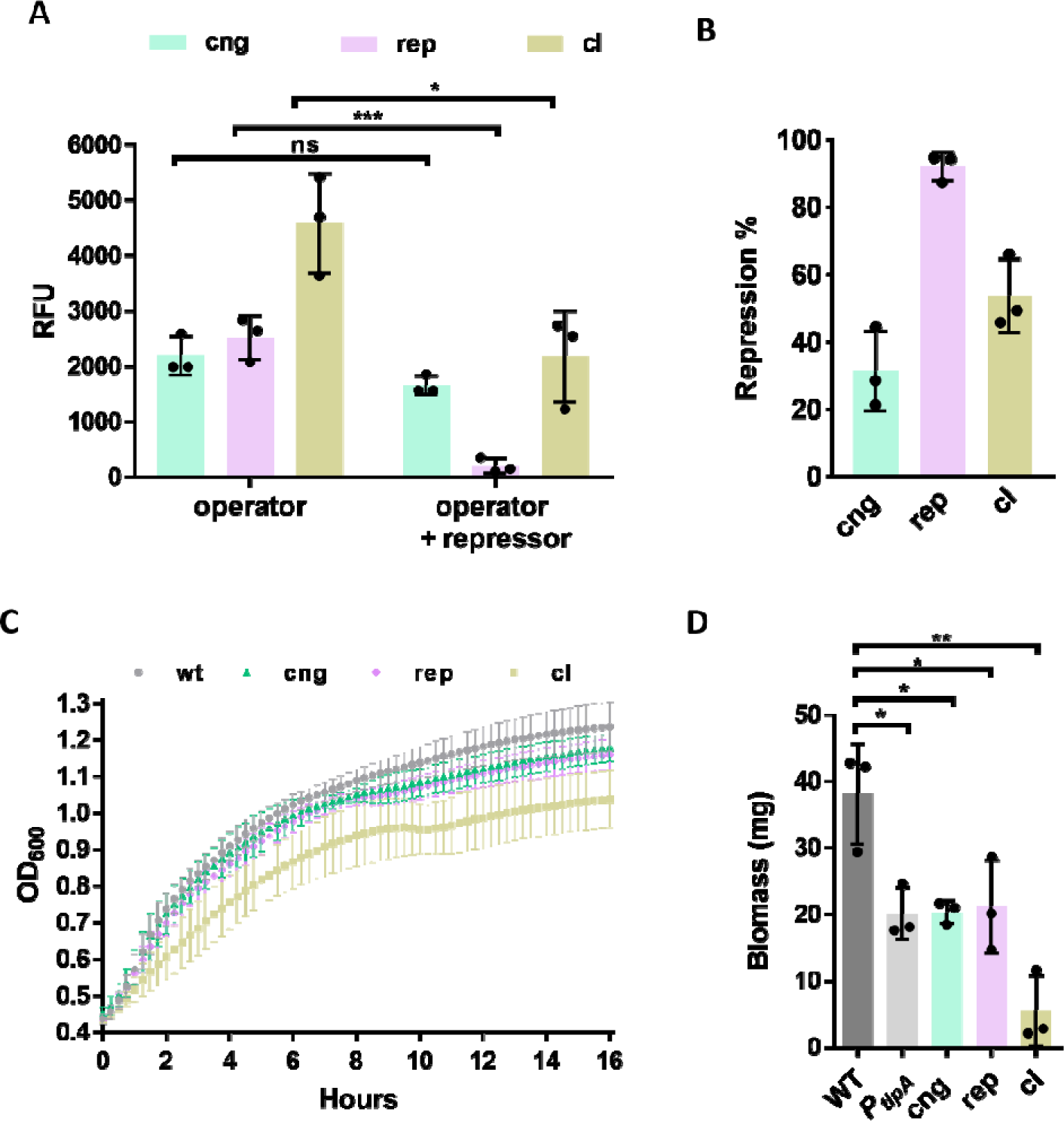
**A)** Expression levels of mCherry in relative fluorescence units (RFU) for all three operators (O cng, O rep and O cI) and all three operators plus repressors (cng, rep and cI). **B)** Percentage of repression mediated by each repressor. **C)** Growth curves of all three repressors and wild-type bacteria over 16 hours. **D)** Bacterial biomass of wild-type, *P_tlpA_*_mCherry, and bacteria carrying each repressor after overnight growth in the incubator. Each sample is based on a 5-mL culture. Experiments for Figures A, B and D were performed as experimental triplicates (N=3). Experiments for Figure C were performed as experimental duplicates, each with two technical replicates (N=2, n=2). Column heights and error bars represent the means and standard deviations (SD). ns = not significant, * *p* < 0.05, ** *p* < 0.01, *** *p* < 0.001.

**Figure 4:**
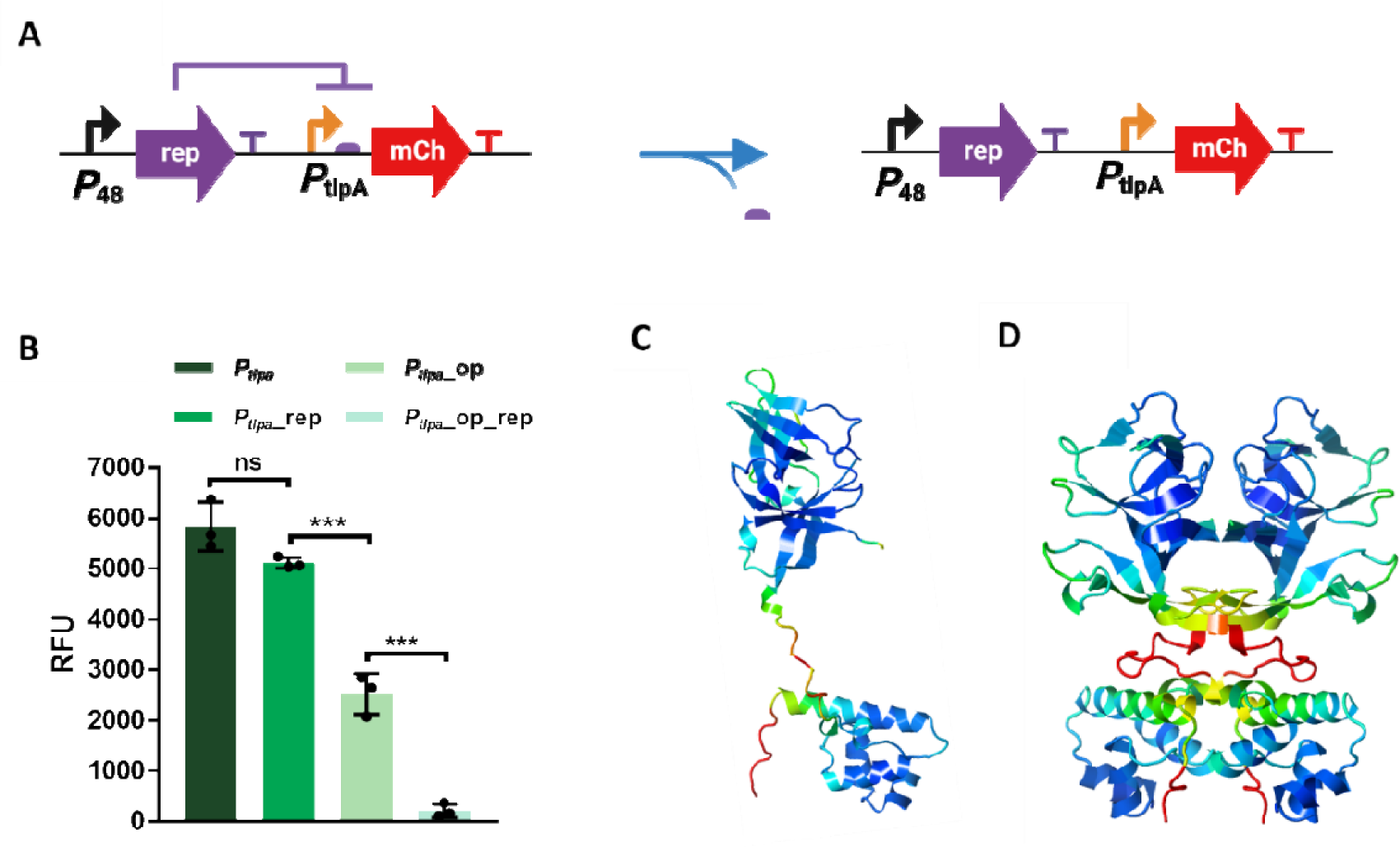
**A)** Scheme of the site-directed mutagenesis done to remove the operator from the plasmid. **B)** Expression levels of mCherry in terms of RFU for *P_tlpA_* (unmodified), *P_tlpA_*_rep (only repressor), *P_tlpA_*_op (only rep operator) and *P_tlpA_*_op_rep (operator plus rep). All the experiments were performed as experimental triplicates (N = 3). Column heights and error bars represent the means and SD. ns = not significant, \*\*\**p* < 0.001. **C)** AlphaFold 3D structure prediction of the rep as a monomer. **D)** AlphaFold 3D structure prediction of the rep as a dimer.

### The operator is essential for repression

We proceeded to confirm that rep repression was dependent on the presence of the operator sequence. This sequence is generally required for the DNA-binding domain (DBD) of the repressors to bind and favor repression. For that, we removed the operator from the plasmid by site-directed mutagenesis PCR using the bacterial pellet as a template for the amplification (**Figure 3A**). The PCR product was then cloned through direct cloning into *L. plantarum* WCFS1 (**Figure S5**). After removing the operator, mCherry expression increased drastically owing to both the inability of rep to bind to repress and the excision of the operator from *P_tlpA_* (**Figure 3B**), which was known to have a negative impact on the expression (**Figure S2**). We also employed Interpro to confirm the presence of a DBD based on a helix-turn-helix domain at the N-terminal of the protein. In addition, AlphaFold predicted the 3D structure of rep, which showed to have the typical structure of a repressor, a DBD at the N-terminal and a dimerization domain at the C-terminal (**Figure 3C**). AlphaFold also predicted the protein-protein interaction of rep monomers forming a dimer with the dimerization domains and DBDs from one monomer associated with those from the other monomer, which is typically needed for repression (**Figure 3D**).

### Optimization of rep-based repression

Next, we attempted to increase the levels of repression by introducing certain modifications to the promoter region. The first approach involved *P_tlpA_* engineering by introducing modifications in the placement of the operator. Thus, three new variants were cloned: i) removing 14-bp between the operator and the RBS (O1), ii) placing the operator within *P_tlpA_*, between the -35 and -10 boxes (O2), and iii) adding an additional operator upstream of *P_tlpA_* (O3) (**Figure 5A**). Removing part of the spacer between the operator and the RBS and placing the operator within *P_tlpA_* considerably decreased the expression of mCherry, even in the absence of the rep (**Figure 5B**). On the other hand, adding an extra operator sequence upstream of *P_tlpA_*only mildly affected mCherry expression but significantly improved repression, from 92% (initial clone, **Figure 3B**) to above 99% (**Figure 5C**). We tested this double-operator approach for the cng repressor by adding an extra operator (O cng) by PCR upstream the *P_tlpA_* promoter(**Figure S6A**) However, surprisingly no repression could be detected after the addition of an extra operator in this repression module **(Figure S6B**). This suggests that even though this double-operator approach is a valid strategy to improve repressor performance, it is not generally applicable to all repressors.

**Figure 5:**
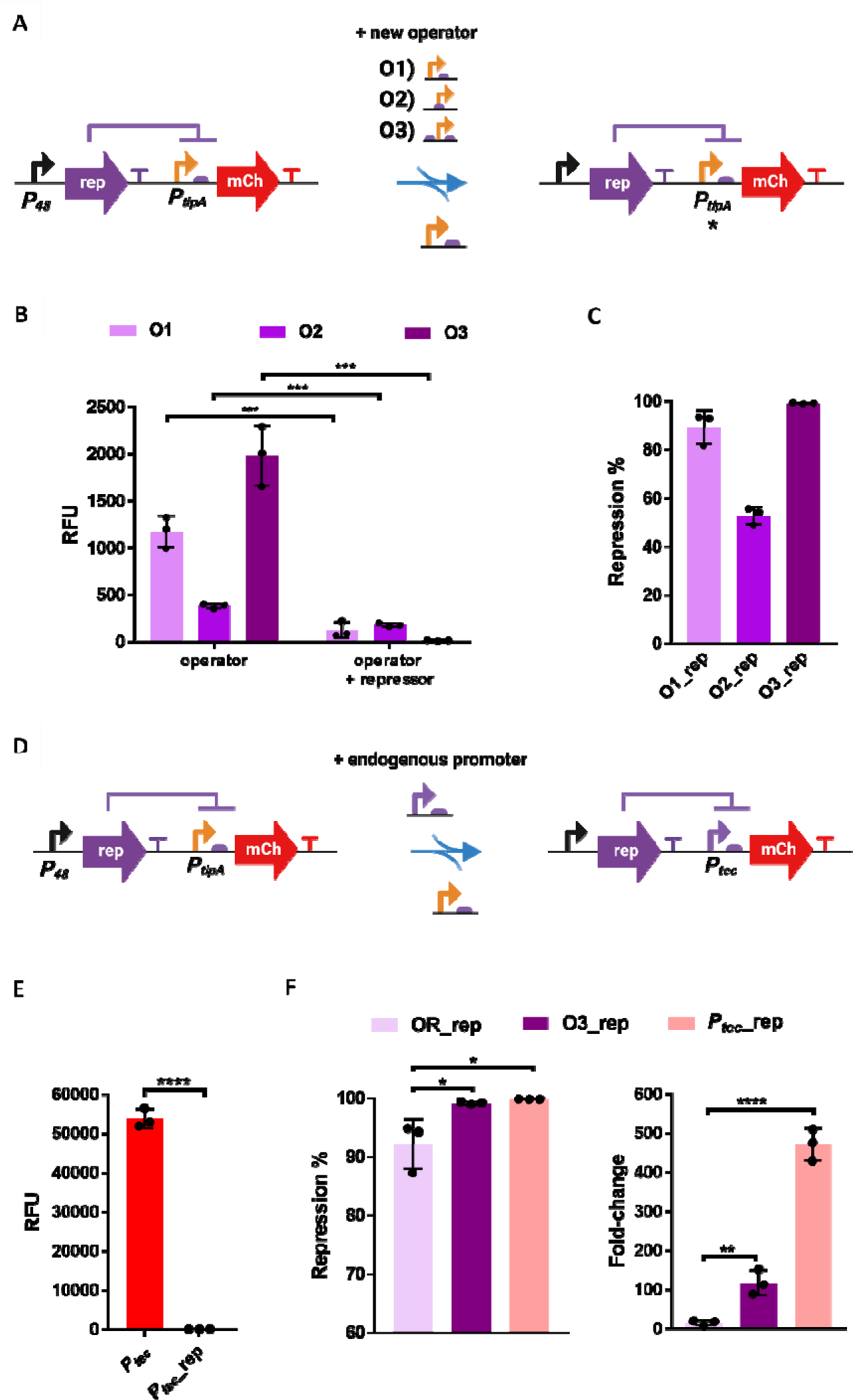
**A)** Scheme of the genetic modifications that introduced the operator of rep at different locations in and around the promoter *P_tlpA_*. **B)** Expression levels of mCherry for all three operator variants (with and without the rep repressor in the plasmid). **C)** Percentage of repression mediated by each operator variant (O1, O2 and O3). **D)** Scheme of the substitution of *P_tlpA_* by *P_tec_***. E)** Expression levels of mCherry for *P_tec_*_mCherry and *P_tec_*_rep_mCherry bacteria. **F)** Percentage of repression and fold-changes for *P_tlpA_*_O_rep (OR_rep), *P_tlpA_*_O3_rep (O3_rep) and *P_tec_* _rep. All the experiments were performed as experimental triplicates (N = 3). Column heights and error bars represent the means and SD. * *p* < 0.05, ** *p* < 0.01, *** *p* < 0.001, **** p < 0.0001.

The second approach attempted to improve repression by replacing *P_tlpA_*with the native promoter (*P_tec_*) associated with rep from the phage mv4 (**Figure 5D**) [36]. In this cloning, we first replaced the rep-operated *P_tlpA_* with *P_tec_* in the plasmid encoding for rep and repression was almost complete. Surprisingly, when the repressor was removed from the plasmid by site-directed mutagenesis (**Figure S7**), the level of mCherry expression was extremely high, due to which the repression efficiency was determined as >99.7% (**Figure 5F** and **5E**).

We also evaluated the effect of these new clones (*P_tlpA_*_O3_rep and *P_tec_* _rep) on cell growth. It was observed that growth was only slightly decreased compared to wild-type bacteria (**Figure S8**), similar to that of rep with a single operator and the cng repressor, and better than that of the cI repressor (**Figure 3C**).

These results proved that repression can be enhanced by either engineering the operator placement within *P_tlpA_* or by introducing the endogenous *P_tec_* promoter. Whereas the original clone showed a fold-change of ∼15 (*P_tlpA_*_O_rep), the optimized clones showed a fold-change of ∼117 (*P_tlpA_*_O3_rep_mCherry) and ∼475 (*P_tec_*_rep) (**Figure 5G**).

### Characterization of a strong constitutive promoter

Next, owing to the surprisingly high expression levels of *P_tec_*, we further characterized it and compared it to *P_tlpA_*. After overnight growth at 37 °C, we could observe a change in the MRS media color due to high levels of mCherry production. This color change was more pronounced when growing these bacteria in LB media supplemented with glucose to favor bacterial growth in this non-optimal media. Such visible color change of the medium due to the expression of a fluorescent protein is regularly observed when expressing these proteins in *E. coli* but has not been reported in lactobacilli. Also, when spinning these bacteria down, the pellet was bright red, similar to what is observed in *E. coli* (**Figure 6A**). As expected, such a strong constitutive promoter had an effect on cell growth compared to wild-type bacteria. This effect was evident when bacteria were grown overnight in the incubator with shaking and not in the plate reader (**Figure S9**). However, biomass was higher than bacteria carrying *P_tlpA__*mCherry, cng repressor and rep repressor.

**Figure 6:**
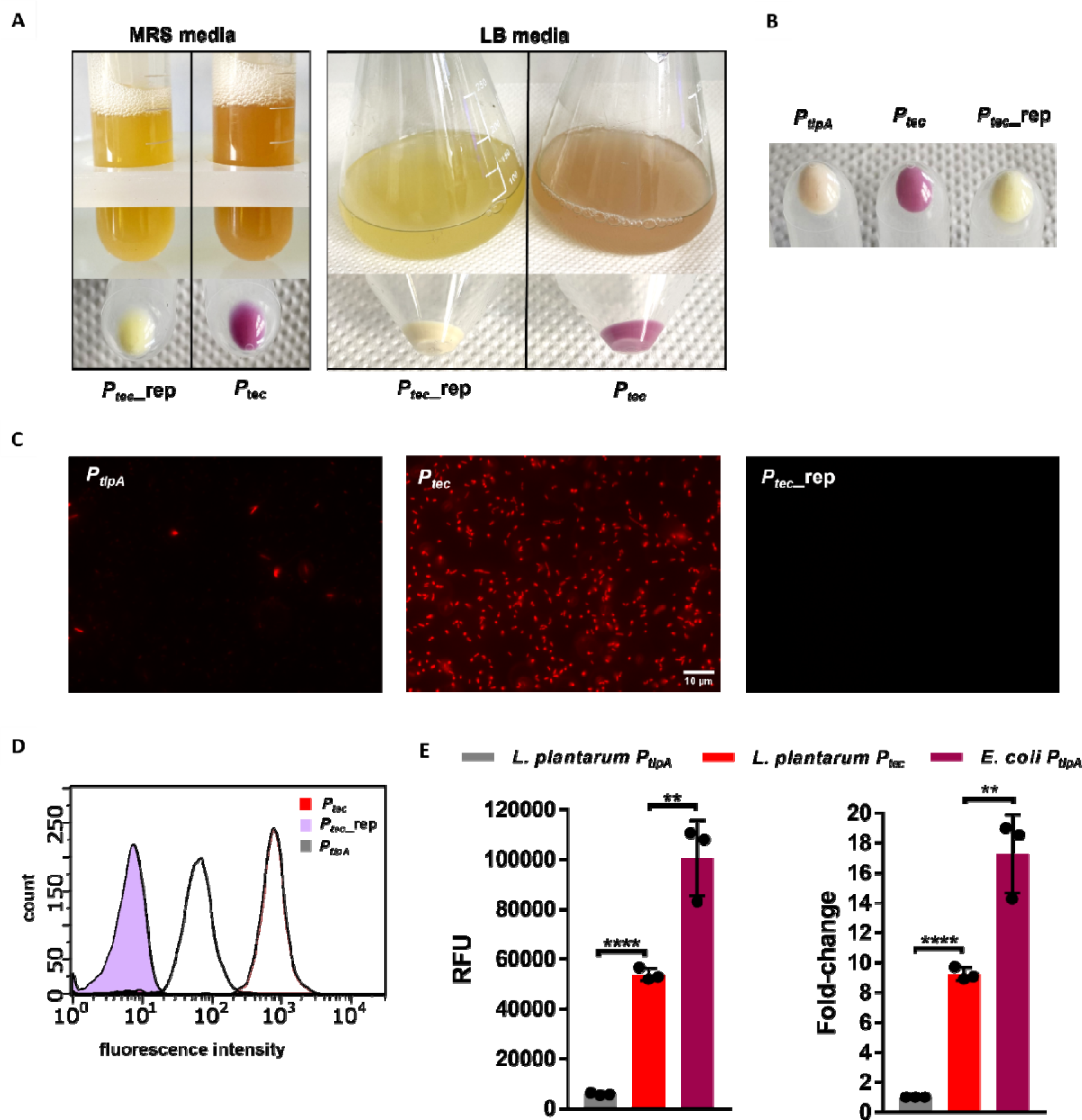
**A)** Left: 5-ml MRS cultures based on *P_tec_*_rep and *P_tec_* bacteria. Cells were pelleted down, and pellets are shown at the bottom. Right: 95-ml LB (supplemented with 5 ml 1M glucose) cultures based on *P_tec_*_rep and *P_tec_* bacteria. Cells were pelleted down, and the corresponding pellets are shown at the bottom. **B)** Pellets of bacteria encoded to express mCherry driven by *P_tec_*_rep, *P_tec_* and *P_tlpA_* after overnight growth. **C)** Fluorescence microscopy images of bacteria encoded to express mCherry driven by *P_tec_*_rep, *P_tec_* and *P_tlpA_*. **D)** FACS data showing the comparison between bacteria encoded to express mCherry driven by *P_tec_* _rep, *P_tec_* and *P_tlpA_*. **E)** Expression levels of mCherry driven by *P_tec_*in *L. plantarum* WCSF1, *P_tlpA_* in *L. plantarum* WCSF1 and *P_tlpA_* in *E. coli* Nissle 1917. Fold changes of mCherry expression driven by *P_tec_* in *L. plantarum* WCSF1 and *P_tlpA_* in *E. coli* Nissle normalized to the expression in *P_tlpA_ L. plantarum* WCSF1. All the experiments were performed as experimental triplicates (N = 3). Column heights and error bars represent the means and SD. ** *p* < 0.01, *** *p* < 0.001.

In comparison with *P_tlpA_*, *P_tec_* drove a considerably higher level of gene expression, which was visible by eye (**Figure 6B**) and microscopy, (**Figure 6C**) and was confirmed by flow cytometry (**Figure 6D**). Fluorescence spectroscopy using a plate reader revealed that *P_tec_*was ∼9 times stronger than *P_tlpA_* (**Figure 6E**), confirming it to be the strongest constitutive promoter discovered for heterologous gene expression in *L. plantarum* WCFS1. Since the visible color changes of the liquid culture and bacterial pellets were comparable to expression in *E. coli*, we compared *P_tec_*-driven mCherry expression levels in *L. plantarum* WCFS1 with the expression in probiotic *E. coli* Nissle 1917 driven by the strong *P_tlpA_* promoter. After quantifying the expression levels of mCherry with the microplate reader, we observed that while *P_tlpA_* in *E. coli* 1917 was 17-fold stronger than in *L. plantarum*, *P_tlpA_* in *E. coli* was only 2-fold stronger than *P_tec_* in *L. plantarum* (**Figure 6E**).

In light of this finding, we attempted to test the strength of the promoter, *P_cpg_*, associated with the 2^nd^ best repressor (cng) identified in this study. Yet, this promoter was only of moderate strength comparable to constitutive promoters like *P_48_* in this strain as it was ∼6 fold weaker than *P_tlpA_* and ∼55 fold weaker than *P_tec_* (**Figure S10**).

## DISCUSSION

Bacteriophages have proven to be an extensive and diverse source of genetic parts to expand the synthetic biology toolbox of bacteria [40], including transcriptional systems [41], integrases [42], anti-CRISPR proteins [43], endolysins [44], and repressors [45]. Such parts are naturally adapted to their bacterial hosts due to the coevolution and the arms race between bacteriophages and bacteria for millions of years [46]. However, bacteriophage parts remain largely unexplored for building genetic circuits in lactobacilli. In this study, we have established a genetic platform for testing transcriptional repressors from lactobacilli-infecting bacteriophages. Relying on operating the strong constitutive *P_tlpA_* promoter proved to be a prudent strategy since the strength and compatibility of the natural phage promoters are unpredictable. For example, *P_cpg_*, the natural promoter associated with the partially effective cng repressor, only showed a moderate expression in *L. plantarum* WCFS1, which might not have been enough to assess repression. Out of six different repressors encoded in genetic switches in lactobacilli prophages, only the rep repressor showed promising results in terms of efficient and reliable repression of the reporter gene without impacting bacterial growth. These results highlight that lactobacilli prophages can be a promising yet challenging source of genetic parts to expand the genetic toolbox. Furthermore, the endogenous promoter associated with rep exhibited unprecedented levels of gene expression, significantly narrowing the gap in terms of expression levels between model (*E. coli* Nissle 1917) and non-model probiotic bacteria engineered for therapeutic applications. *P_tec_* could also be operated and used as a genetic platform to identify more repressors in this strain since even low repression could be more readily assessed than with *P_tlpA_*. The highly efficient *P_tec_*/rep promoter/repressor system could now be applied in combination with other genetic parts for building genetic circuits. One limitation of the system is that it is currently not inducible. However, this provides the opportunity for future work to employ repressor engineering strategies and modify rep into a switchable repressor that responds to sugars [47] or physical stimuli such as light [48] or heat [49]. Such switchable repressors will enable inducible gene expression, which would be desirable to circumvent the notable metabolic burden and stress that *P_tec_*might be causing to the cells due to its transcriptional expression strength. Also, the unmodified repressor can be combined with other inducible gene expression systems to invert the induction system as a NOT logic gate [50, 51]. In combination with the *P_tec_* promoter, such induction or inversion functions can be achieved at a remarkably high level of performance.

## CONCLUSIONS

We identified a novel, strong and efficient promoter/repressor system in the probiotic bacteria *L. plantarum* WCFS1 by screening for such genetic parts in lactobacilli-infecting bacteriophages. After improving the system, we achieved repression levels of >99% and fold-changes of >100. Moreover, we discovered a super strong constitutive promoter, *P_tec_*, which can drive levels of expression never achieved before in this strain, precisely ∼9 times higher than the previously reported strongest promoter, *P_tlpA_*. These novel genetic parts will be instrumental in expanding the capabilities to engineer gene expression regulation in *L. plantarum*.

## LIST OF ABBREVIATIONS

LAB: Lactic Acid Bacteria
MRS: De Man, Rogosa and Sharpe.
PCR: Polymerase Chain Reaction.
rpm: Revolutions per minute.
RFU: Relative Fluorescent Units.
RBS: Ribosome Binding Sites.
OD_600_: Optical density at 600 nm.
SD: Standard deviations.

## SUPPLEMENTARY INFORMATION

The online version contains supplementary material available at DOI: XXX. All relevant sequences for this study have been included in **Figure S11**.

## DECLARATIONS

### Ethics approval and consent to participate

Not applicable.

### Consent for publication

Not applicable

### Availability of data and materials

The datasets used and/or analysed during the current study are available from the corresponding author on reasonable request.

### Competing interests

A patent application has been filed based on the results of this work (Application no. is DE 102022 119024.2).

### Funding

This work was supported by a Deutsche Forschungsgemeinschaft (DFG) Research grant [Project # 455063657 - https://gepris.dfg.de/gepris/projekt/455063657] and the Leibniz Science Campus Living Therapeutic Materials.

### Authors’ contributions

MBA and SS developed the initial concept. MBA conducted most of the experiments and wrote the manuscript, partially assisted by AW. VST conducted the experiments related to *E. coli* Nissle 1917 engineering and analysis. SS supervised the study and revised the manuscript. All authors have read and approved the final manuscript.

## Acknowledgements

The *Lactiplantibacillus plantarum* WCFS1 strain was a kind gift from Prof. Gregor Fuhrmann (Helmholtz Institute for Pharmaceutical Research, Saarland). The plasmid pLp_3050sNuc was a kind gift from Prof. Geir Mathiesen (Addgene plasmid # 122030). *E. coli* Nissle was a kind gift from Prof. Rolf Müller. *L. plantarum* Lg1e was a kind gift from Dr. Makiko Kakikawa. Biorender was used for generating the schemes for the figures of this publication.

