## Supporting Information for "Discovery of a high-performance phage-derived promoter/repressor system for probiotic lactobacillus engineering"

| **REPRESSOR** | **PHAGE** | **HOST** | | **SEQUENCE** | **OPERATOR** |
| --- | --- | --- | --- | --- | --- |
| **cpg** | øg1e | *Lactobacillus plantarum* | MASSGIGNRLKELRNMQGKTQDEVAKSIGISRARYSHLENERNEPDNELLKLLASYYEVSTDYLLGNSEKSHKSPDWATEADRIDLDKLLQSNTPMGYGGMSMAPEDKEKVRNVIEGIYWDRLKKLREEGKK | | GATACGATATGTATC |
| **cng** | øg1e | *Lactobacillus plantarum* | MKRERLIAERNRNGWSQNSVAKLLDIAEITVRSIENGSRNPSSKLIAKFSYLFEVKPEILFPDIFLPDKDTKRIISAKANKLTKEAAK | | GATACACAATGTATC |
| **rep** | mv4 | *Lactobacillus delbrueckii* | MPRANYTPQEKQLKSVIAGNLNALLSKTAYKKADVVRQTGISESTVYDYFNGKVLPSPKNVEKLADFFRVSNEEIDPRFATMPENMVPVDQSHLVKIPLIGHIACGEPITADQNIEGYITEYFPDHVDPDSIFALKCEGDSMEPYILDGDIAYIRQQPEVEDGEIAAVLVDGDTRASLKRVKKVGNQVFLLPDNPHYSPIVLDQDHPGKIIGKMIKMSRFQ | | CCCGGTCTAGAACGGGG |
| **tec** | mv4 | *Lactobacillus delbrueckii* | MPKISVRAARVNAGFSQDEAAKKLGISRFTLQRYESDPKQIRQGMLEKMRLVYNMDHDNLFFKI | | ACAAAAAGTCAACTTTTTTCAACTTTTTGT |
| **cI** | A2 | *Lactobacillus casei* | MKTNDEIIKTLNDLRNREGISISELARRVDMAKSSVSRYFNGTREFPLNYVDKFASALHTTPESLIGVSPVDPFKVKKLNVHSYPYIPAEISAGILCNVDPLTSDDVETIQLPDSVMGRYAGDSSILMMHINGESMNQTIPDGSLIAVKQYNDIQDLKDGDIVVFADDGDYAVKYFYNDRQKQIVTFIPDSTDKRFSPIMYTYEDLEEENIKIIGRVVVYTVVL | | CCCAAAAGGAAACGAAAGGA |
| **cro** | A2 | *Lactobacillus casei* | MGTRMIFFRCSQKETKGGNPMTLNLKRLRAERIAKGMNQDEMAKAMGWHTRSSYAKRENGITTISATELVKMASILGYGTNQLDLFFTNNVPDRERKGMTV | | CCCTATTGTGCACAATCGGG |

**Table S1:** List of all the repressors included in this study. The origin, host, sequence, and corresponding operator are indicated per repressor.

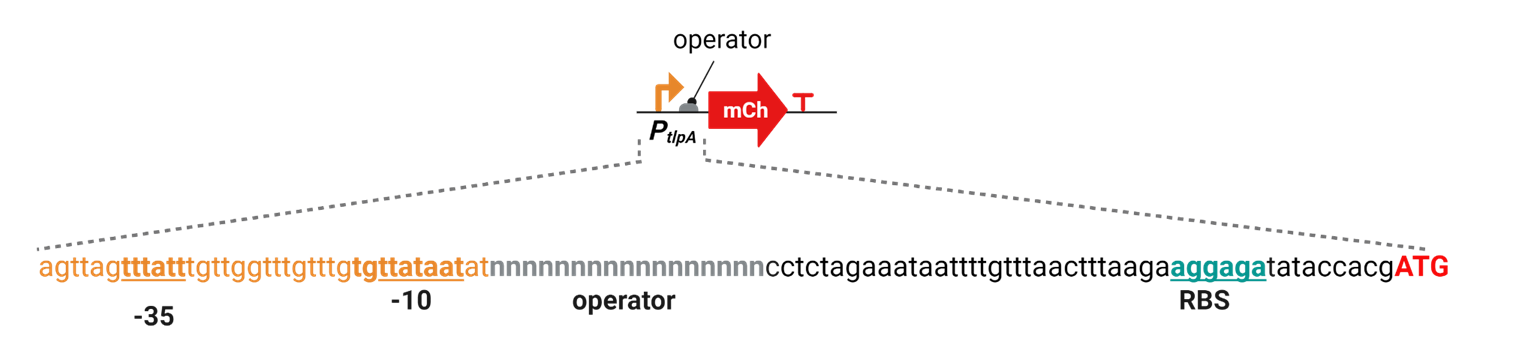

**Figure S1:** Placement of the operator in the promoter region. Each operator sequence was placed downstream *P_tlpA_*.

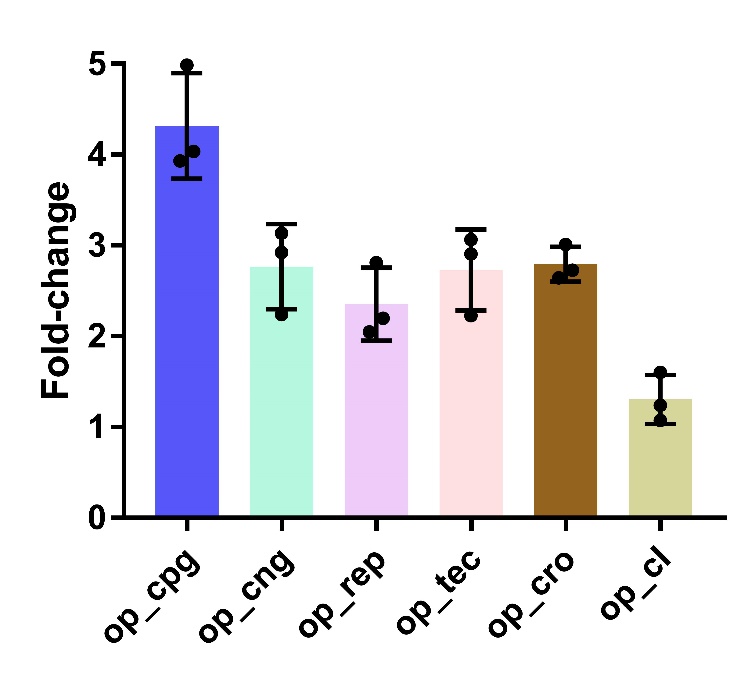

**Figure S2:** Fold-changes of *P_tlpA_* (unmodified promoter) compared to *P_tlpA_* operated with operators from each repressor tested in this study. This indicates the effect of placing each operator in *P_tlpA_*. All the experiments were performed as experimental triplicates (N = 3). Column heights and error bars represent the means and SD.

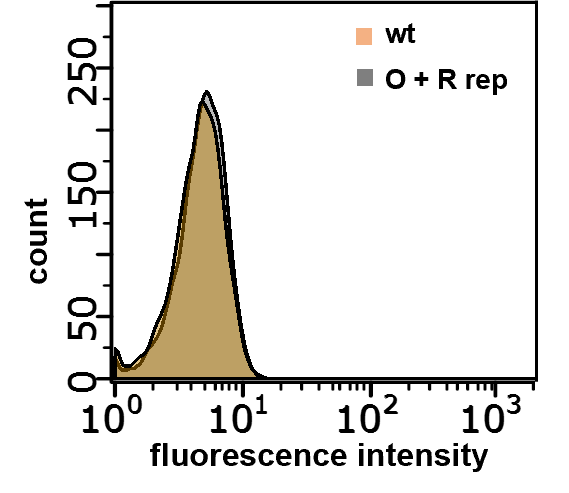

**Figure S3:** FACS data showing the comparison between bacteria carrying the rep repressor (repressing mCherry) and wild-type bacteria.

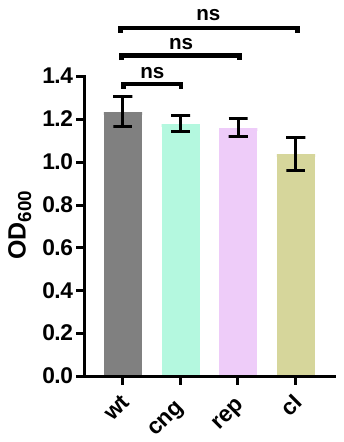

**Figure S4:** OD_600_ values after 16 hours of growth in the microplate reader for wild-type bacteria and bacteria encoding for each repressor. Column heights and error bars represent the means and SD. Ns = not significant.

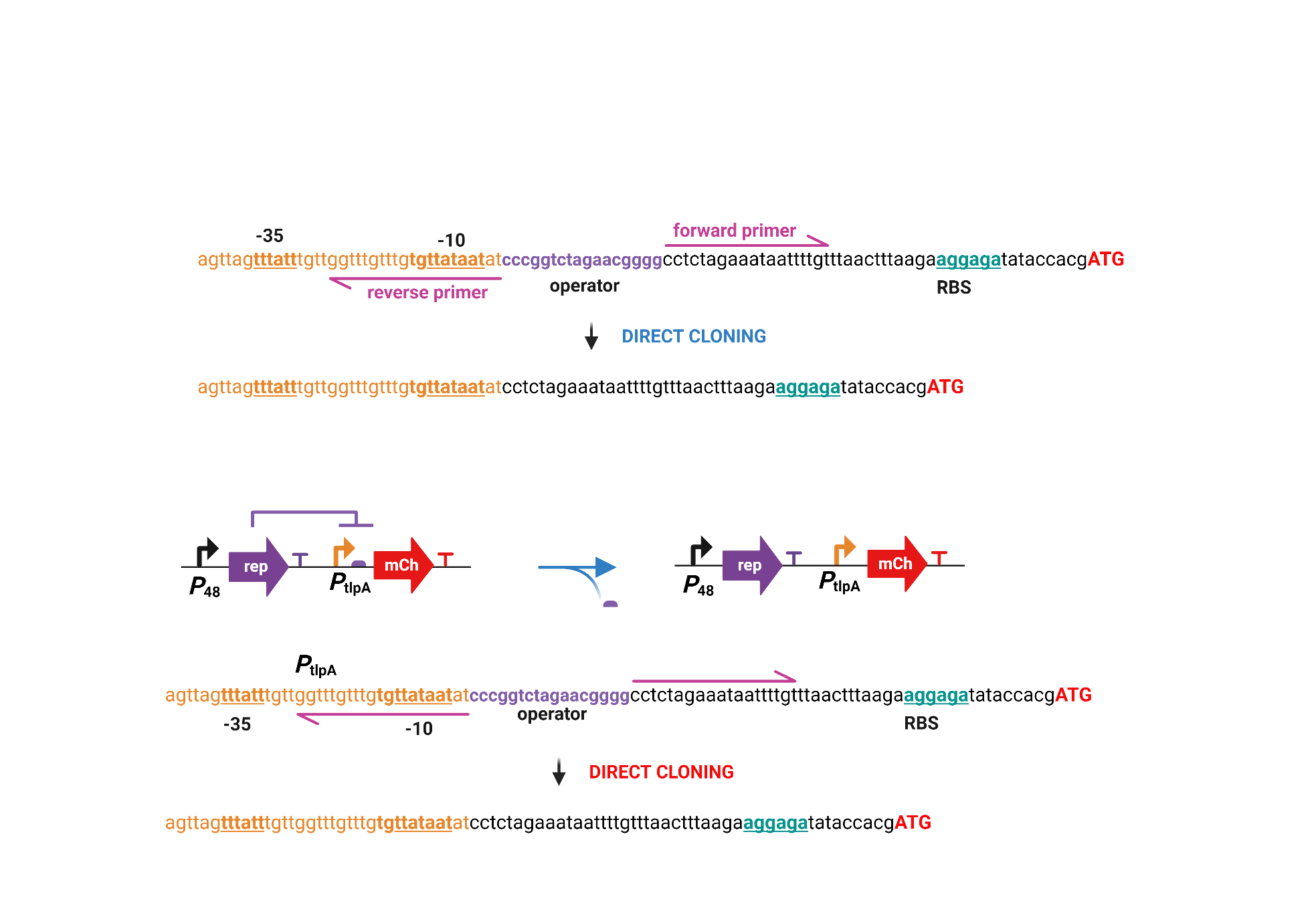

**Figure S5: A)** Scheme of the site-directed mutagenesis PCR to remove the operator sequence from the promoter region of the plasmid.

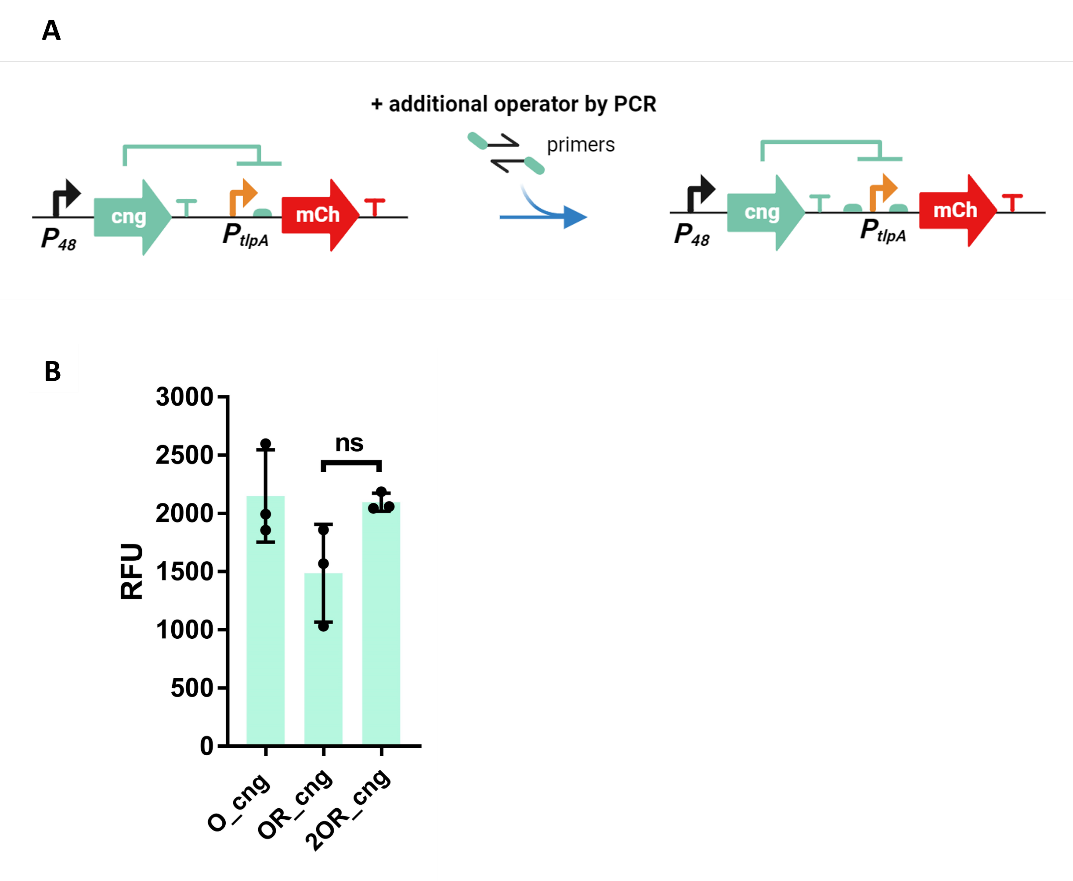

**Figure S6: A)** Scheme of the additional PCR to insert an additional operator sequence upstream the promoter. **B)** Expression levels of mCherry in terms of RFU for the clones based on single cng operator (O_cng), cng operator plus repressor (OR_cng) and double cng operators plus repressor (2OR_cng). All the experiments were performed as experimental triplicates (N = 3). Column heights and error bars represent the means and SD. Ns = not significant.

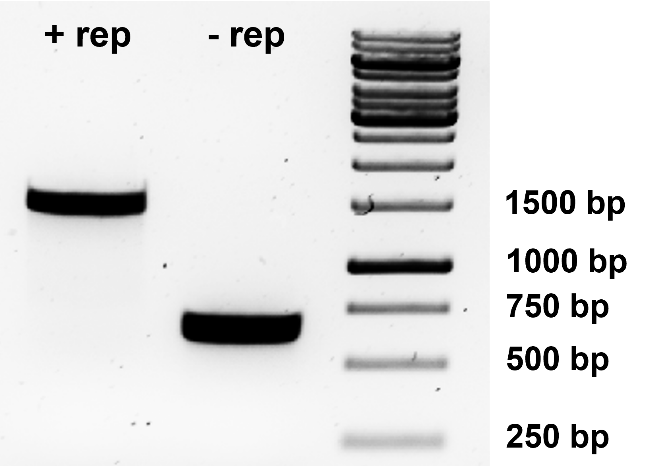

**Figure S7:** Agarose gel showing that the excision of rep from the plasmid by site-directed mutagenesis took place as expected. The “+ rep” product (1524 bp) is 891 bp bigger than the “- rep” product (633 bp) due to the excision of the *P_48_*_rep gene. Bacterial pellets were used as a template for the PCR. Generuler 1 Kb DNA Ladder was used for the reference.

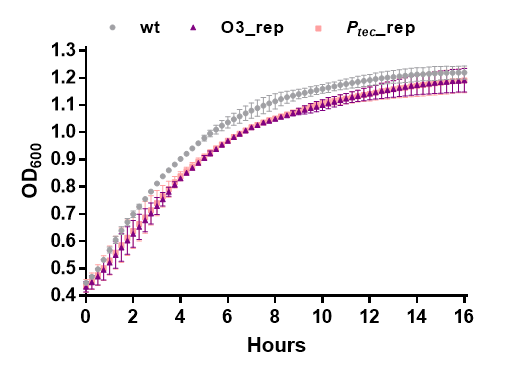

**Figure S8:** Growth curves of *P_tlpA_*_O3_rep_mCherry_,_ *P_tec_* _rep_mCherry and wild-type bacteria over 16 hours. Experiments were performed as experimental duplicates, each with two technical replicates (N=2, n=2). Data points and error bars represent the means and SD.

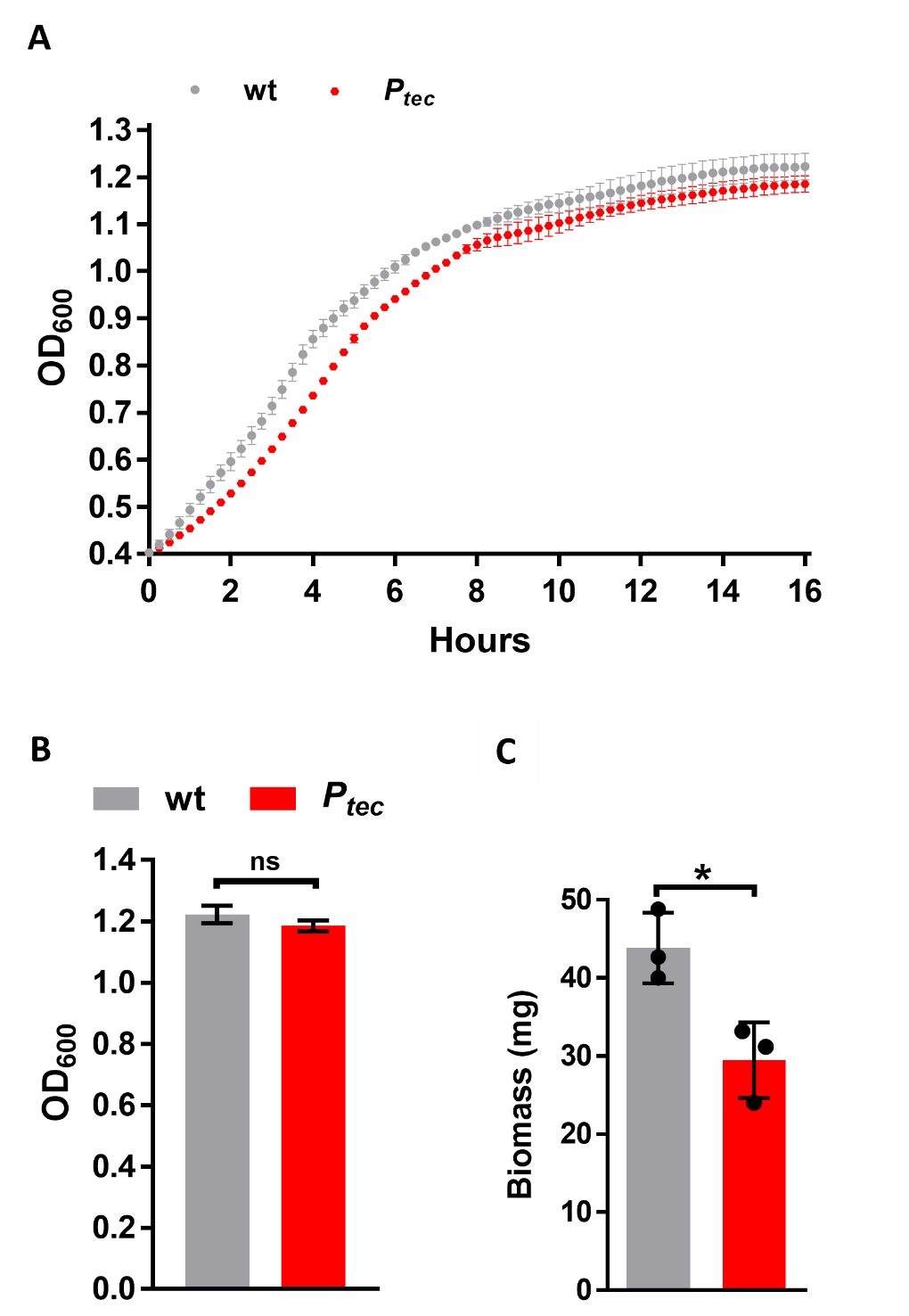

**Figure S9: A)** Growth curves of *P_tec_* _mCherry and wild-type bacteria after 16 hours of growth in the microplate reader. **B)** OD_600_ values for *P_tec_* _mCherry and wild-type bacteria 16 hours of growth in the microplate reader. **C)** Bacterial biomass of *P_tec_* _mCherry and wild-type bacteria after overnight growth in the incubator. Each sample is based on a 5-mL culture. Experiments for Figures A and B were performed as experimenta**l** duplicates, each with two technical replicates (N=2, n=2). Experiments for Figure C were performed as experimental triplicates (N=3). Symbols or column heights and error bars represent the means and SD. ns = not significant, * *p* < 0.05.

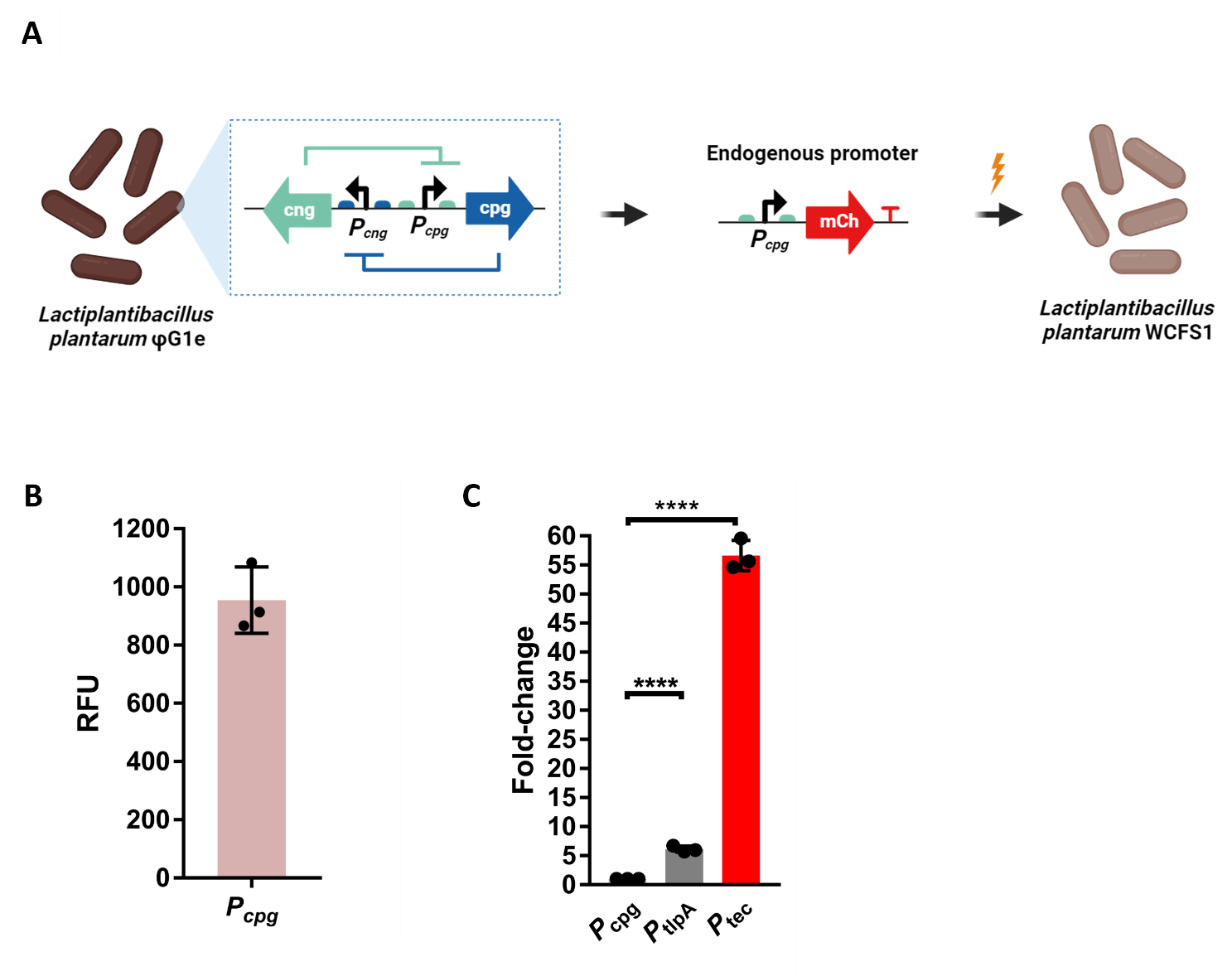

**Figure S10: A)** Scheme of the *P_cpg_* cloning. The promoter was amplified from the genome of *L. plantarum* ϕg1e and inserted in the mCherry-encoding plasmid. **B)** Expression levels of mCherry in terms of RFU for the *P_cpg__*mCherry bacteria. **C)** Fold changes between *P_cpg_* and *P_tlpA_* and *P_tec_* promoters in *L. plantarum* WCFS1. All the experiments were performed in triplicate. Column heights and error bars represent the means and SD.

**“O1” sequence:** *Modified clone, one operator sequence downstream P_tlpa_, shorter spacer.*

agttagtttatttgttggtttgtttgtgttataatat**cccggtctagaacgggg**tttgtttaactttaaga**aggaga**tataccacg**atg**

**“O2” sequence:** *Modified clone, one operator sequence within P_tlpa._*

agttagtttatt**cccggtctagaacgggg**ttgttataatatcctctagaaataattttgtttaactttaaga**aggaga**tataccacg**atg**

**“O3” sequence:** *Modified clone, two operators, upstream and downstream P_tlpa._*

**cccggtctagaacgggg**ttagtttatttgttggtttgtttgtgttataatat**cccggtctagaacgggg**cctctagaaataattttgtttaactttaaga**aggaga**tataccacg**atg**

***“P*_tec_” sequence:** *Different promoter, endogenous P_tec_ promoter from L. delbrueckii*

tttgtgttgact**cccggtctagaacgggg**tattattaaagcatcaagaacgaa**aggaga**aaacgaa**atg**

**Figure S11:** Relevant sequences generated in this study. *P*_tlpA_ promoter is shown in orange. *P*_tec_ promoter is shown in violet. Rep operator is shown in violet and bold. The -35 and -10 boxes are underlined. The RBS sequence is shown in blue and in bold. The mCherry start codon is shown in red and bold.

| **NAME** | **SEQUENCE (5’-3’)** |
| --- | --- |
| **Operator insertion fw** | atggtttcaaagggtgaagaag |
| **Operator insertion rev** | ttacgatgcagatctccg |
| **Repressor insertion fw** | ctttctgcgctgcattaagccggtagatcagcatagatct |
| **Repressor insertion rev** | ttacgatgcagatctccg |
| **HiFi PCR repressors fw** | catatctgacgacttagc |
| **HiFi PCR repressors rev** | gtctatgctctattacgat |
| **Rep operator excision fw** | cctctagaaataattttgt |
| **Rep operator excision rev** | atattataacacaaacaaacc |
| **Rep optimization fw** | atggtttcaaagggtgaa |
| **Rep optimization rev** | tctatctatatctctagcgag |
| **HiFi PCR rep optimization fw** | ccgttagcgtagtagtagc |
| **HiFi PCR rep optimization rev** | cttcaaagggtcaacagc |
| **Sequencing GOI fw** | cgttactaaagggaatggag |
| **Sequencing GOI rev** | agtggaacgaaaactcac |
| **Cng + operator fw** | ataatatctagtttatttgttggtttg |
| **Cng + operator rev** | atgtatcactaacaaacaaattaaacc |
| **Cng ϕg1e fw** | tactaaagggaatggagaccccaactaaatcagcatgactc |
| **Cng ϕg1e rev** | ttcttcaccctttgaaaccatagtaccgctccttt |
| ***P_cpg_* fw** | ttttgcacctccaattcc |
| ***P_cpg_* rev** | ctagtagttgagattgcca |

**Table S2:** List of all the primers used in this study.
